# Screening of a pooled library of chimeric antigen receptor T cells based on secretory function

**DOI:** 10.1101/2025.07.02.662376

**Authors:** Citradewi Soemardy, Anna Mei, Rocio Castellanos-Rueda, Nikol Garcia Espinoza, Monika Kizerwetter, Jamie B Spangler, Sai T Reddy, Dino Di Carlo

**Affiliations:** Department of Bioengineering, University of California Los Angeles, Los Angeles, California 90095, USA; Department of Biosystems Science and Engineering, ETH Zürich, 4056 Basel, Switzerland; Life Science Zurich Graduate School, ETH Zürich, University of Zurich, 8057 Zürich, Switzerland; Department of Chemical and Biomolecular Engineering, Johns Hopkins University, Baltimore, Maryland 21231, USA; Department of Biomedical Engineering, Johns Hopkins University, Baltimore, Maryland 21231, USA; Jonsson Comprehensive Cancer Center, University of California Los Angeles, Los Angeles, California 90095, United States; California NanoSystems Institute (CNSI), University of California Los Angeles, Los Angeles, California 90095, USA

## Abstract

Chimeric antigen receptor (CAR) T cell therapies have shown promise in treating hematologic malignancies, but challenges remain due to immune suppression, antigen heterogeneity, and insufficient functional screening platforms. Here, we present a modular nanovial-based platform for high-throughput, single-cell functional screening of pooled CAR T cell libraries. Nanovials, hydrogel microparticles with nanoliter-scale cavities, were functionalized with recombinant HER2 antigen and cytokine-capture antibodies to simulate antigen-presenting cells and capture secreted interferon-γ (IFNγ). This system enabled the selective capture, activation, and functional profiling of CAR T cells based on antigen engagement and cytokine secretion. We screened a 32-variant CAR library with diverse intracellular signaling domains, using nanovials to isolate IFNγ-secreting cells after 3- and 12-hour CAR-specific stimulation. IL15RA-containing CARs, particularly IL15RA-CD28, were preferentially enriched in the sorted T cells after 3 hours of stimulation, consistent with early effector activation profiles. By 12 hours, IL15RA-containing constructs remained enriched while other CD40-containing domains showed delayed but substantial enrichment, suggesting prolonged signaling dynamics. The platform’s high-throughput capability (>2 million cells screened), compatibility with downstream sequencing, and tunable antigen presentation make it ideal for identifying CAR constructs associated with various time-dependent secretion phenotypes.

## Introduction

Immunotherapy has emerged as a key pillar of cancer treatment alongside surgery, radiotherapy and chemotherapy and harnesses the immune cells of the patient to combat cancer. Successful approaches of immunotherapies range from enhancing endogenous immune cell activity through checkpoint inhibition to employing genetically engineered immune cells, such as chimeric antigen receptor (CAR) T cells, to specifically target and kill cancer cells. While CAR T cell therapies have demonstrated remarkable success in achieving long-lasting complete remission in patients with hematologic malignancies^1^, significant challenges persist while treating solid tumors. These include immunosuppression within the tumor microenvironment, poor immune cell infiltration, and tumor antigen escape. In contrast, checkpoint inhibition and engineered T cell receptor (TCR) therapies have shown clinical efficacy in treating solid tumors such as carcinomas, melanoma and synovial cell sarcoma^2,3^. One possible explanation for the observed discrepancy might be that CARs and engineered TCRs engage different signaling pathways which can lead to the secretion of different effector molecules^4^. This could result in the variation of their effectiveness when targeting solid tumors^5^.

Conventional screening methods to evaluate CAR functionality only analyze a handful of diverse CAR T cell constructs in parallel. In these screens, potent CARs are mainly selected based on functional phenotypes such as their antigen affinity, antigen specificity, cytotoxicity and cytokine production profiles. Because characterizing individual CAR variants is a very time- and labor-intensive process, only a few different constructs can be produced and tested at the same time. In order to facilitate the discovery of novel CAR T cell treatments and to expand the space of tested CAR domains, multiple research labs have developed various high-throughput screens combined with computational analysis in recent years^6^. In these studies, different strategies to screen for the single-chain variable fragment (scFv) or intracellular domains (ICDs) in CAR constructs were established to identify more effective CAR designs^7–11^. After long-term pooled cytotoxicity assays with antigen-expressing target cells, potent CAR T cells were selected based on their activation, proliferation, and cytokine production. These methods deepen our understanding of how a certain CAR design can impact T cell behavior and shape CAR T cells towards a desired phenotype. However, these methods still rely on bulk co-culture of a pooled CAR T cell library with target cells, which obscures cooperative effects between T cells containing different constructs. As such, there remains the need for single-cell technology that allows the phenotypic screening of pooled CAR libraries based on their binding, activation and cytokine-secretion profile while capturing the functional differences, such as variations in cytokine secretion, of individual CAR constructs within a pooled library. Additionally, there is not yet a system that allows tunable control of antigen levels on target cells to enable selection and screening of CAR T cells based on nuanced differences in antigen sensitivity, which is critical for reducing on-target, off-tumor effects.

Microfluidic technologies that encapsulate single cells into microfluidic droplets, optoelectronic pens, and microwell arrays have been applied to measure secretions from single cells^12^. Droplet-based assays enable measurement of cytokine secretions that accumulate to high concentrations in the small droplet volume and allow for downstream sorting with custom droplet sorters^13–15^. However, measuring both CAR T cell affinity to a target antigen and its triggered activation requires a solid phase or other cell that is expressing the target antigen. Microwell array techniques where target-expressing cells and CAR T cells are loaded together can measure secretions triggered by antigen engagement and CAR clustering but remains a low-throughput manual process^16–18^. Optoelectronic tweezer-based systems, such as the Beacon system from Bruker Cellular Analysis^11,19,20^, can sort cells from single pens after characterizing T cell secretory function or killing of target cells, but this system is limited to screening less than ten thousand cells with significant equipment cost.

In contrast, flow cytometry-based methods can screen large CAR T cell libraries in high-throughput^7,11^. Due to its wide-spread availability and versatility, flow cytometry is commonly used to evaluate surface markers of activation on CAR T cells and cytokine secretion profiles through intracellular cytokine staining (ICS). Using flow cytometry-based methods, Goodman et al. screened a pooled CAR T cell library in multiple rounds of antigen stimulation assays, followed by the quantification of higher levels of CD69, a marker of activation, intracellular cytokine (IFNγ and IL-2) production, and proliferation^7,9^. Unfortunately, ICS detects only cytokines accumulated within the cell and therefore may not be correlated with the time-dependent secretion of cytokines or other secreted effector molecules that remain stored in granules before activation, like granzyme B.

Our lab has previously developed a high-throughput single-cell functional screening technology using hydrogel microparticles containing a nanoliter-scale cavity, termed nanovials (*fig1*)^21,22^. Nanovials have been previously applied for discovery of TCR repertoires against various viral and tumor-specific peptides and linking secretory phenotype to surface marker expression in both human and murine T cells^23,24^. This platform does not require complex microfluidic handling and is compatible with standard cell culture techniques and a range of downstream analysis tools including fluorescence activated cell sorting (FACS) and single-cell RNA sequencing (scRNA-seq)^21–23,25–28^.

**Figure 1.**
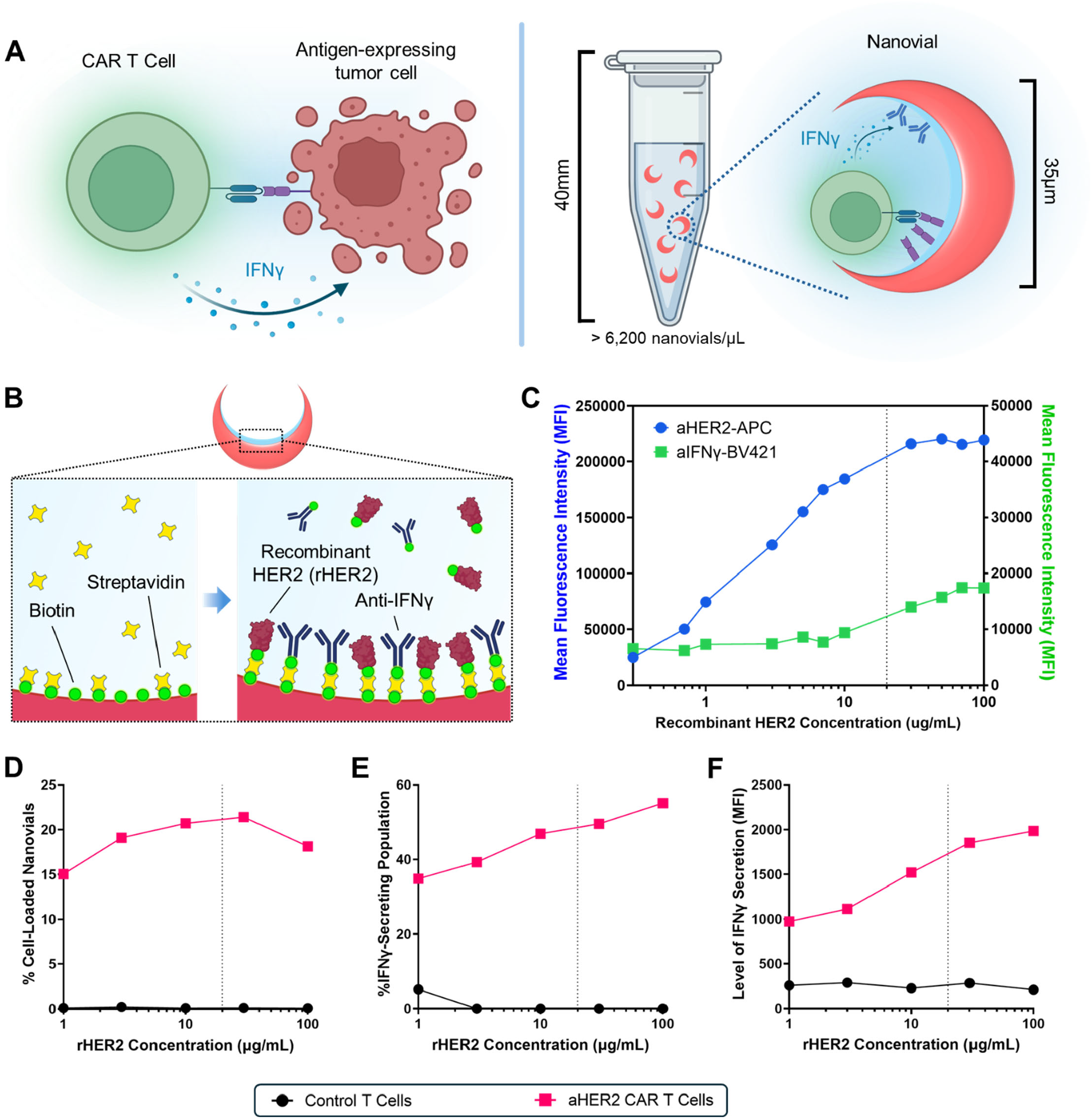
(A) Schematic of engagement of a CAR T cell with a tumor cell expressing the target antigen, leading to cytokine release. An antigen-conjugated nanovial can mimic the antigen-expressing tumor cell, leading to selective capture and activation of the CAR T cell and release of cytokines. (B) Schematics showing the cavities of nanovials functionalized using biotin-streptavidin chemistry and subsequently conjugated with recombinant HER2 (rHER2) antigens and antibodies against IFNγ to capture and quantify the secretion level of a bound cell. (C) Mean fluorescence intensity (MFI) signals from a population of nanovials functionalized with varying concentrations of rHER2 and aIFNγ antibody at 20μg/mL concentration are plotted. To measure the ability to capture cytokines across rHER2 concentrations, nanovials were also incubated with 50ng/mL recombinant IFNγ before labeling. Nanovials were stained with aHER2-APC antibody and aIFNγ-BV421 detection antibody (D) For a cell to nanovial ratio of 1.6:1, the percentage of nanovials loaded with cells is plotted for control T cells (circles) and aHER2 CAR T cells (squares) as a function of rHER2 levels on the nanovials. (E) Percentage of IFNγ-secreting fraction from the cell-loaded nanovials is plotted after 3 hours of incubation as a function of rHER2 levels on the nanovials. (F) The MFI of IFNγ signal on nanovials loaded with control T cells and aHER2 CAR T cells is plotted after 3 hours of incubation as a function of rHER2 levels on the nanovials.

Here, we functionalized nanovials with recombinant human epidermal growth factor 2 (rHER2) antigens to serve as artificial tumor associate antigen (TAA)-displaying cells (*Fig. 1A*), achieving the selective capture of anti-HER2 (aHER2) CAR T cells based on antigen binding, and targeted CAR T cell activation to induce the secretion of IFNγ. Secreted IFNγ was subsequently captured on the nanovials and associated CAR constructs could be sorted and sequenced. We were able to characterize the antigen-density dependent activity of the CAR T cells and applied this system to screen for a library of CARs transfected into T cells with different combinations of ICDs, including CD28, 4-1BB, CD40, IL15RA, and CTLA4. Over 2,000,000 cells were screened from the library in a single run resulting in 300-500 sorted cells based on high secretion function. IFNγ secreting and non-secreting CAR T cells were sequenced to identify constructs from the library with an enhanced cytokine secretion profile. CAR variants containing IL15RA were enriched when measuring secreted IFNγ accumulated over 3 hours, which persisted in secretion measurements over 12 hours. This result reflected other findings highlighting IL-15-based signaling as critical in promoting an effector phenotype in T cells, resulting in release of effector molecules, including IFNγ^29–31^. Overall, the nanovial platform provides unique functional data to screen and characterize large libraries of CARs to select for variants with desired phenotypes that may be more efficacious in solid tumors.

## Results

### Functionalization of nanovials to screen CAR T cell libraries

In order to develop a system capable of screening libraries of CAR T cells based on cytokine secretion, we first functionalized the nanovial surfaces with (i) the tumor-associated antigen (TAA) HER2 to facilitate capture and activation of CAR T cells, and (ii) antibodies to enable detection of secreted cytokines. We used a microfluidic device to generate uniformly sized hydrogel microparticles containing nanoliter-scale cavities using aqueous two-phase separation of polyethylene glycol (PEG) and gelatin in generated microdroplets, following previously described approaches *(Fig. S1A)*^23^. The nanovials were measured to have an average outer diameter of 34μm and average cavity diameter of 22μm, sufficient for single cell loading. Leveraging the lysine residues in the gelatin layer lining the nanovial cavity, we conjugated biotin to the surface of the nanovial cavity to allow modular modification of nanovials *(Fig. 1B, Fig. S1A)*. For selective capture and stimulation of aHER2 CAR T cells, we functionalized nanovials with recombinant HER2 (rHER2) antigen. Nanovials were simultaneously functionalized with cytokine capture antibodies against interferon-γ (aIFNγ) (*Fig. 1B*), to detect secreted IFNγ. As both modification steps use shared biotin sites on the nanovial, we first evaluated the dynamic range of cytokine detection when conjugating the nanovials with multiple types of antibodies simultaneously. Previous experiments using nanovials conjugated with four different antibodies, including an aIFNγ capture antibody, at 1:1:1:1 ratio (130nM each) showed a fluorescence intensity dynamic range spanning over two orders of magnitude when detecting up to 1000ng/mL recombinant IFNγ (*Fig. S1B*). This enabled detection of cytokine concentrations as low as 10ng/mL. We decided to focus on IFNγ, a widely-recognized pro-inflammatory cytokine released by activated T cells to mediate immune responses^32–34^ that is also used in CAR-T cell potency assays.

We next created nanovials with tunable amounts of rHER2 antigen, simulating cells with different HER2 expression levels. We functionalized nanovials in a solution with 20μg/mL (130nM) aIFNγ capture antibody and different concentrations of rHER2 antigen, ranging between 0 and 100μg/mL. Maximal HER2 levels were reached between 10-30μg/mL of rHER2 (*Fig. 1C, Fig. S2*). Additionally, we incubated the nanovials with 50ng/mL recombinant IFNγ to assess whether cytokine capture efficiency was maintained across these different ratios of rHER2 to aIFNγ. IFNγ capture levels remained relatively constant, independent of the concentration of rHER2 used (*Fig. 1C, Fig. S2*).

### Selective capture and activation of CAR T cells using nanovials functionalized with recombinant HER2 antigens

Using rHER2-functionalized nanovials, we hypothesized that we could selectively capture and activate aHER2 2^nd^ generation CD28-CD3Z CAR T cells (28Z CAR), leading to the accumulation of detectable IFNγ secretion within the nanovial cavity. In order to evaluate the amount of antigen needed for selective CAR T cell binding and activation, we first varied the concentration of rHER2 conjugated onto the nanovials (1-100μg/mL). Interestingly, the percentage of cell-loaded nanovials did not change significantly across this range (*Fig. 1D*), leading to stable binding to rHER2 on nanovials. However, we observed a dose-dependent increase of the fraction of IFNγ-secreting cells as rHER2 antigen concentration increased (*Fig. 1E*). Similarly, we observed a 2-fold increase in the mean fluorescence intensity (MFI) associated with increased IFNγ secretion levels across this antigen range (*Fig. 1F)*. Based on these findings, we selected 20μg/mL (270nM) as the default rHER2 antigen concentration for subsequent experiments.

To evaluate the selectivity of cell capture and activation, untransfected human T cells, used as control T cells, or CAR T cells were incubated for 3h with either aCD45- or rHER2-functionalized nanovials (*Fig. 2A,B-Left*). Following this incubation, nanovials were assessed for cell loading and IFNγ secretion. We previously showed that aCD45-conjugated nanovials effectively capture T cells from human peripheral blood mononuclear cells (PBMCs) without stimulating them^23^, acting as a negative control. Previous studies have also shown that the level of cell loading into nanovials follows a Poisson distribution^22,23,25^. As such, we adjusted the cell-to-nanovial ratio to 1.6:1 to target a 10-20% cell loading efficiency while minimizing instances of multiple cells being loaded into a single nanovial. aCD45 nanovials captured both control T cells (∼7% cell-loaded nanovials) and 28Z aHER2 CAR T cells (∼12% cell-loaded nanovials) (*Fig. 2A,B-Mid*). In contrast, rHER2 nanovials selectively captured 28Z aHER2 CAR T cells (∼8% cell-loaded nanovials). Gating on these cell-loaded nanovials, we observed IFNγ secretion for aHER2 CAR T cells loaded into rHER2 nanovials by flow cytometry, confirming antigen-specific activation (*Fig. 2A,B-Right*). These results were confirmed by observation of live CAR T cells (green) and their IFNγ accumulation (blue) within the nanovial cavity using fluorescence microscopy as shown in *Fig. 2C*.

**Figure 2.**
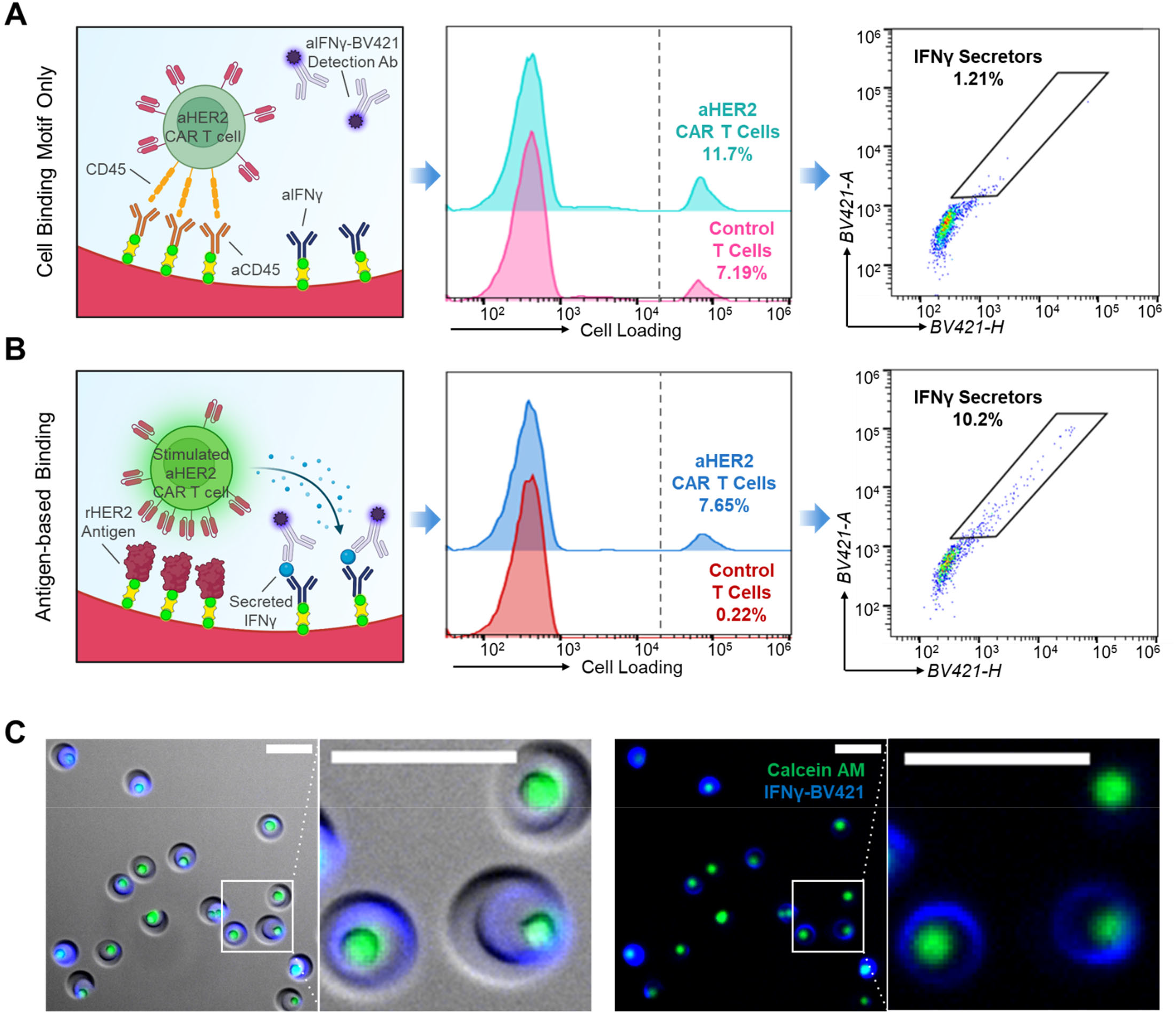
Selective binding and secretion of aHER2 CAR T cells. (A) Nanovial functionalization with aCD45, cell loading histograms, and IFNγ scatter plots showing binding of CAR T cells and basal levels of IFNγ secretion. Both aHER2 CAR T cells and control T cells can be captured in aCD45-conjugated nanovials. (B) Nanovial functionalization with rHER2 antigen, cell loading histograms, and IFNγ scatter plots showing selective binding of CAR T cells and increased fraction of secreting cells and levels of IFNγ secretion. (C) Fluorescence microscopy images of the sorted IFNγ-secreting population from the gate in B with (left) and without (right) brightfield overlay. Cells are stained green with calcein AM viability dye. Secreted IFNγ, bound to the nanovials, is labeled in blue. Scale bar: 50µm.

We also evaluated the time dependence of IFNγ accumulation within the nanovial cavity to determine an optimal secretion time point. Incubation times longer than 6 hours, up to 12 hours, led to overall increase in IFNγ signal on cell-containing nanovials, but also led to a rise of cross-talk signal–defined as IFNγ signal detected in empty nanovials (*Fig. S3*). Additionally, prolonged incubation resulted in a reduction of the fraction of nanovials loaded with cells, presumably due to enhanced motility or increased receptor endocytosis of activated CAR T cells (*Fig. S4*). In contrast, incubation time between 0 and 6 hours showed minimal differences in both cell-loading efficiency and cross-talk levels (*Fig. S5*), while still producing detectable IFNγ signals. We therefore selected a 3-hour incubation period for our initial screen of a CAR library.

### Sequencing of enriched CAR constructs from a pooled library identified IL15RA-containing variants as IFNγ secretors, indicating their role in enhancing cytokine secretion and potential therapeutic efficacy

We next applied the platform to screen a library of aHER2 CAR T cells with varying combinations of intracellular domains (ICDs). The pooled library is composed of 32 variants, resulting from recombining the ICDs of 5 different immune cell receptors (CD28, 41BB, IL-15 receptor α (IL15RA), CD40, and CTLA4) into 1^st^ generation, 2^nd^ generation, or 3^rd^ generation CAR constructs, all of which include CD3Z domain (*Fig. 3A*)^9^. In addition, a non-signaling (NS)-CAR construct lacking any ICD and CD3Z domain serves as an internal negative control allowing cells to bind to their target antigen without potential CAR-dependent activation. CD28 and 41BB represent commonly used costimulatory domains in many CAR T cells studies, including clinically approved CAR T cell products^1,35–39^. IL15RA and CD40 signaling domains have shown enhanced antitumor capabilities in preclinical studies^29,40^. Lastly, despite being known as an inhibitory receptor of T cells, CTLA4 has shown improved antitumor capability when incorporated into CAR constructs^41^.

**Figure 3.**
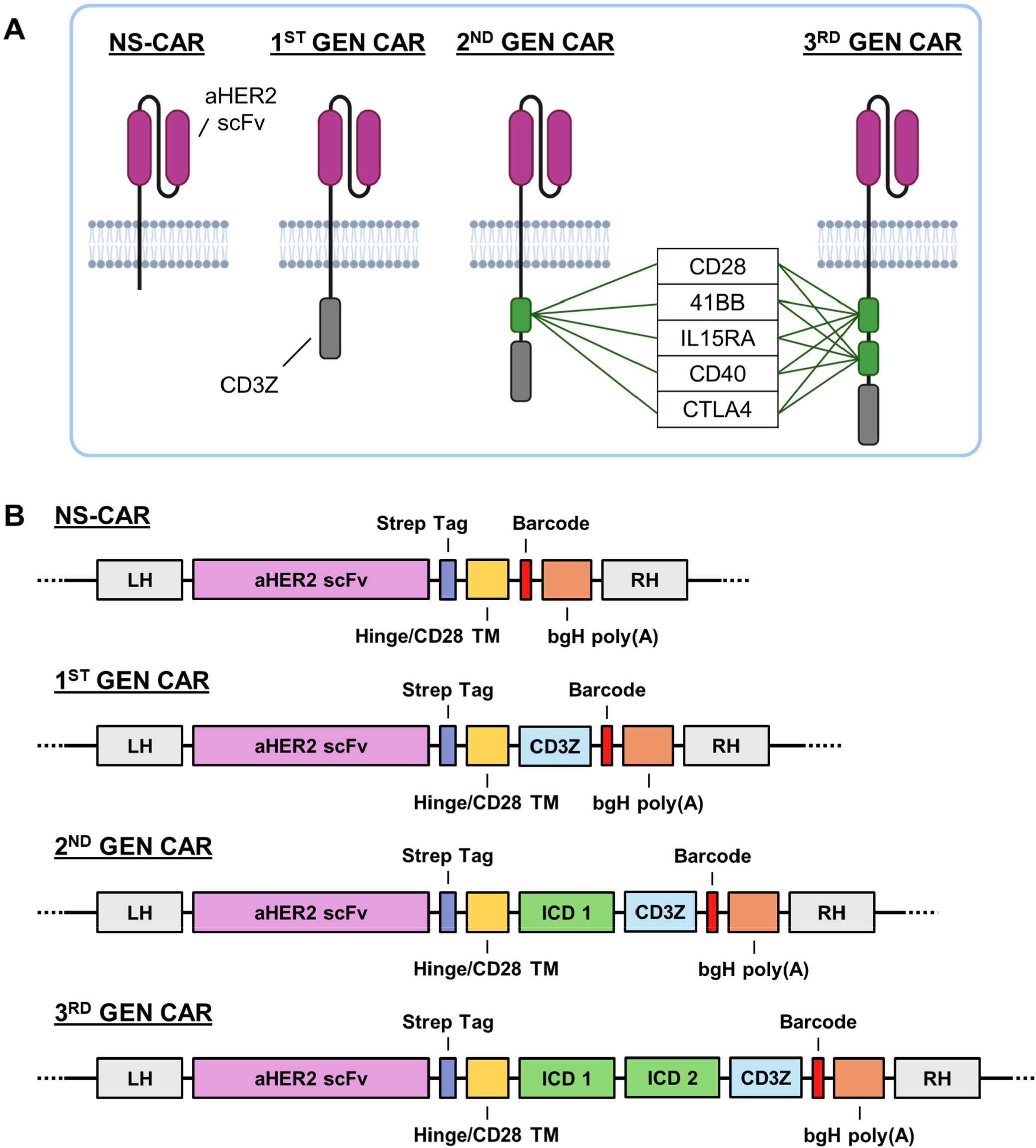
CAR library construction. (A) Design of the 32-variant aHER2 CAR T cell library. The library includes (n=1) non-signaling, NS-CAR, (n=1) 1^st^ generation CAR containing only CD3Z domain, (n=5) 2^nd^ generation CARs, and (n=25) 3^rd^ generation CARs with varying mixtures of ICDs. (B) The CAR transgene designs are shown which is inserted by CRISPR-Cas9 and HDR into the 1^st^ exon of the T cell receptor alpha constant (TRAC) locus. LH – left homology; RH – right homology; TM – transmembrane domain.

The CAR library was genomically integrated into the TCR alpha constant (TRAC) locus of primary human T cells using CRISPR-Cas9 and homology-directed repair (HDR). This method allows for controlled and precise insertion of the CAR gene, while knocking out the endogenous TCR^42^ (*Fig. 3B*). In order to identify the individual CAR constructs in the library after the IFNγ secretion assay, each CAR variant contained a unique DNA barcode sequence. We confirmed the successful enrichment of the CAR library and of the standard aHER2 2^nd^ generation CD28-CD3Z CAR construct (28Z CAR) after FACS through Flow cytometry (*Fig. S6*). When sequencing the library, we found that each of the construct barcodes was represented to varying amounts, although the fraction of each construct was not uniform. (*Fig. S7, Table S5*).

Library CAR T cells were loaded into nanovials functionalized with rHER2 antigen and aIFNγ capture antibodies and incubated for 3 hours to allow accumulation of secreted IFNγ, enabling evaluation of differences in secretion levels among the CAR variants. Under the same loading conditions as the 28Z CAR construct, where we observed 8-15% of nanovial loading, only 2–3% of nanovials were loaded with library CAR T cells. Lower binding of the library variants to the nanovial might be due to effects of the intracellular domains (ICDs), as the scFv was the same across the entire library and it is known that ICDs have an influence on the CAR expression levels and receptor endocytosis^43^. To study the pro-inflammatory potential of the different CAR candidates, we classified cell-loaded nanovials based on IFNγ secretion: those with IFNγ signal above a basal threshold (determined using aCD45 control nanovials) were designated as “IFNγ secretors,” and those below the threshold were designated as “Non-secretors.” Using FACS, we sorted nanovials positively gated for IFNγ secretion (*Fig. 4A*). ∼12,000 “Non-secretors” events and ∼300-500 “IFNγ secretors” events were sorted and processed for sequencing.

**Figure 4.**
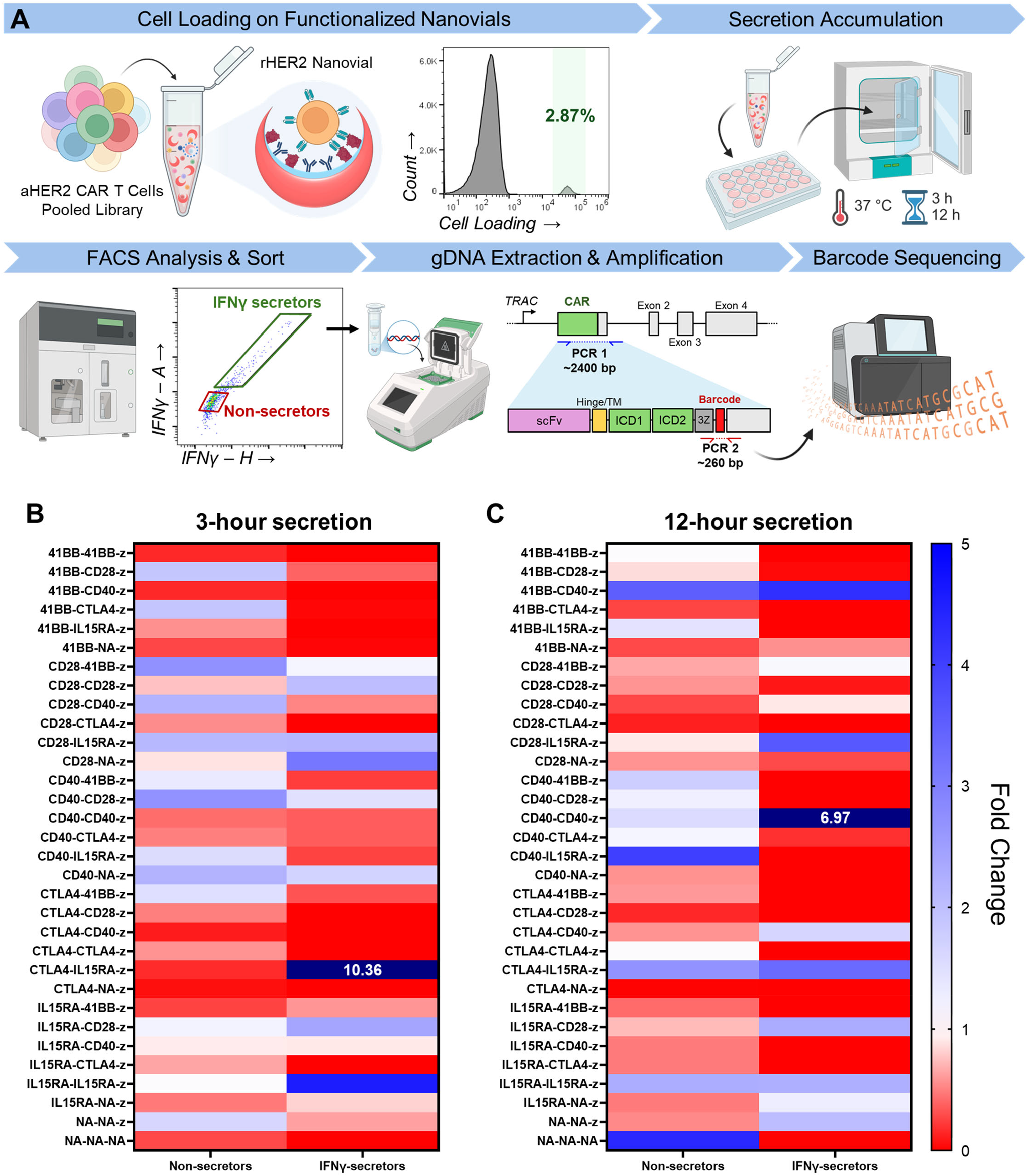
CAR library screening workflow and results identifying constructs enriched based on secretory function. (A) Screening workflow and representative results of CAR T cell loading, flow cytometry histogram and IFNγ scatter plot of secretion signal for the library. Key steps include cell loading on rHER2 functionalized nanovials, secretion accumulation for 3 hours or 12 hours, fluorescence-activated cell sorting (FACS) of cell populations on nanovials based on secretion phenotype, gDNA extraction and amplification, and barcode sequencing and analysis. (B) Table showing relative depletion (values from 0-1) or enrichment (values >1) of constructs normalized to their respective starting library frequency for gated and sorted “Non-secretors” and “IFNγ secretors” after a 3-hour incubation. (C) As for **B** but for a 12-hour incubation.

Following sorting, amplicon sequencing of barcode regions enabled analysis of the selective enrichment and depletion of specific constructs based on binding and secretion of IFNγ (*Fig. 4A, Table S3, Table S4*). For our screen following a 3-hour incubation period, several variants containing the IL15RA signaling domain were preferentially enriched compared to their starting library frequency. These include CTLA4-IL15RA, IL15RA-IL15RA, and IL15RA-CD28 (*Fig. 4B, Table S5*). Here, the first domain corresponds to the membrane-proximal domain, while the second is the membrane-distal domain. The CAR construct containing CTLA4-IL15RA showed the highest level of enrichment compared to the initial library at ∼10-fold. Corroborating this result, CTLA4-IL15RA was also observed to be depleted in the “Non-secretors” population. On the other hand, IL15RA-NA as a single domain 2^nd^ generation CAR construct was not enriched from the starting library, indicating that the presence of IL15RA alone may not be sufficient to drive secretion. However, the IL15RA-IL15RA (double IL15RA) CAR construct showed substantial enrichment in the “IFNγ secretors” population, ∼5-fold higher compared to the initial library. As expected, both negative controls, including the construct without any ICDs (NA-NA-NA) and the construct with only CD3ζ (NA-NA-Z) were depleted in the “IFNγ secretors” population (*Fig. 4B*). The presence of CD3ζ (NA-NA-Z) appeared to facilitate binding to rHER2 leading to relative enrichment in the “Non-secretor” population. In a second run of the 3-hour secretion screen (biological replicate), we again observed enrichment of constructs containing the IL15RA domain (IL15RA-CD28 and IL15RA-CTLA4) despite the library being less uniformly distributed (*Fig. S7A*,*B*). We also found the CD40-NA construct to be enriched in the repeated assay by ∼4-fold.

Overall, our level of confidence in enrichment or depletion in the assay varies across constructs based on the relative abundance of the constructs in the starting library (*Fig. S8A, Table S5*). For example, constructs such as CTLA4-CTLA4, CTLA4-CD40, CTLA4-CD28, CTLA4-41BB, and CTLA4-NA were below 0.30% abundance in the initial library of the first screen while the expected abundance was ∼3%. Other constructs were better represented in the initial library and we can be more confident in our enrichment results, such as for CTLA4-IL15RA, IL15RA-IL15RA, IL15RA-41BB, CD40-IL15RA, CD40-CD40, and depletion of 1^st^ generation CARs (NA-NA-Z) and NS-CARs. Of note, constructs containing CTLA4 tend to be underrepresented in the CAR T cell library which may be because this domain has been previously shown to drive receptor internalization and degradation in resting T cells^44,45^.

To understand if particular ICDs may contribute to longer and sustained secretion of IFNγ, we conducted another screen following a 12-hour incubation period with rHER2-functionalized nanovials (*Fig. S8B*). Compared to the 3-hour timepoint, we found that IL15RA-CD28 constructs continued to be enriched, while other IL15RA-containing constructs (CTLA4-IL15RA and IL15RA-IL15RA) remained enriched in “IFNγ secretors” but became enriched in the “Non-secretors” fraction as well (*Fig. 4C*). Additionally, we also see CD28-IL15RA construct becoming more enriched in the “IFNγ secretors” as compared to the “Non-secretors”. While this may confirm the role of IL15RA in inducing IFNγ secretion, it also shows that activation by IL15RA ICD may lead to increased localization of T cells to nanovials compared to other constructs. On the other hand, some CD40 constructs became enriched in the “IFNγ secretors”, including CD40-CD40 and CTLA4-CD40 constructs, with the CD40-CD40 construct showing the highest enrichment at ∼7-fold. Interestingly, most other constructs containing CD40 (CD40-41BB, CD40-CD28, CD40-CTLA4, and IL15RA-CD40) showed significant depletion, compared to the initial library and the “Non-secretors” fraction. Lastly, we also observed enrichment of 41BB-CD40 construct in both the “IFNγ secretors” and the “Non-secretors” population compared to the initial library distribution. Enrichment in the “Non-secretors” population implies that cells with these constructs bound more stably to the antigen-coated nanovials than cells containing other constructs.

## Discussion

This study demonstrates the application of a modular nanovial-based platform for high-throughput, single-cell functional screening of CAR T cell libraries in response to a target antigen. By isolating CAR T cells that selectively bind to a recombinant antigen in nanoliter-scale hydrogel cavities, we were able to quantitatively assess antigen-specific capture and induction of IFNγ secretion over activation windows ranging from 3 to 12 hours. When screening a library of 32 CAR variants, we observed that the enrichment profile of the pooled library differed between the two timepoints. Constructs containing largely IL15RA were enriched following 3 hours of stimulation, whereas some IL15RA and CD40 constructs showed greater enrichment after 12 hours. These findings align with previous work by Nair et al., which demonstrated that IL15RA containing CAR T cells are primed for rapid Th1 effector responses, characterized by IFNγ and IL-2 secretion as early as 6 hours post-antigen stimulation, compared to more conventional CD28-based CAR T cells^29^. In contrast, the delayed enrichment of the CD40-CD40 construct is consistent with results from Castellanos-Rueda et al., which found that CD40-containing CAR designs elicit sustained growth and enrichment from a pool of CAR T cells following repeated antigen stimulation over 12 days^9^. Our study also shows that the IFNγ secretion profile varies with time, which may indicate constructs that are more involved in early versus late activation, or even in persistence of T cell activation. Interestingly, many of the conventional 2^nd^ and 3^rd^ generation CAR T cell constructs containing CD28 and 4-1BB, with the exception of CD28-NA, are not as enriched using this assay method at 3 or 12 hours compared to the IL15RA-CD28 variant.

We have identified constructs yielding varying levels of induced IFNγ at different time points following activation with rHER2 antigen. Further validation of these constructs in the context of *in vitro* co-culture assays measuring IFNγ secretion, proliferation, and cell killing can better map the nanovial screening results to other established assays, and ultimately *in vivo* studies of CAR T cell anti-tumor efficacy. Improving the distribution of each CAR variant in the starting library and optimizing the nanovial functionalization to enhance the yield of CAR T cell binding and activation can further increase the abundance of each construct assessed in each assay and the statistical robustness of the screen.

The quantitative nature of the nanovial platform can be further leveraged by selecting CAR T cell variants with specified cytokine secretion levels, leading to functional profiling of CAR T cells across a continuum of secretion levels and facilitating the identification of constructs that are not only antigen-sensitive but also functionally potent at defined cytokine levels. Constructs that elicit low, intermediate, or high levels of secreted cytokines could be tested for efficacy and safety differences in co-culture and *in vivo* assays. It may be expected that too high levels of secreted cytokines could lead to other toxicities, such as cytokine release syndrome, or be associated with exhausted T cell phenotypes that do not result in the most durable responses^9,33^.

Our results show that the nanovial platform can assist with the rapid optimization of CAR T cells, enabling researchers to quickly sift through large libraries and pinpoint ideal candidates based on cytokine secretion profiles and further finetune the phenotype of a given CAR variant of interest. This format may provide a distinct advantage over traditional pooled assays that evaluate CAR function through proliferation and activation markers in co-culture with target cancer cells, as it enables the study of early effector dynamics and isolates early effector function of a single cell from confounding variables such as paracrine influences and bystander effects. Our approach focuses primarily on the direct interaction between the CAR T cell and its target antigen, isolating antigen-specific CAR T cell activation from additional co-stimulatory and co-inhibitory signals present on tumor cells. To precisely characterize the effect of other potential signals on CAR T cell function during the screen, nanovials can be functionalized with additional types and levels of ligands to further simulate relevant aspects of tumor cells, e.g., using recombinant proteins or virus-like particle carriers to more naturally present membrane proteins^46^. Directly loading a target tumor cell as bait on the nanovial is a parallel approach to provide all potential signals^47^, although this method reduces the ability to isolate specific factors.

A key advantage of the nanovial platform is that it allows for precise control over antigen density. By tuning the amount of rHER2 presented on the nanovial cavity surface, we may be able to selectively enrich for CAR T cells that respond only to specific antigen thresholds, thus improving tumor selectivity and potentially minimizing on-target off-tumor toxicity. This feature is particularly relevant for tumor-associated antigens (TAAs) like HER2, which are overexpressed in tumor cells but are also present at lower levels in healthy tissues. The platform’s ability to screen for CAR constructs that are activated only at higher TAA densities can help identify candidates with optimal specificity and therapeutic efficacy.

While this study primarily focused on IFNγ secretion as a marker of CAR T cell activation, the nanovial platform is capable of multiplexed cytokine detection. Prior studies in our lab have demonstrated the capability to capture and measure additional cytokines and effector molecules secreted by single cells, such as IL-2, TNFα, and granzyme B^23,24^, which provide further insights into the polyfunctional nature of CAR T cells. This multiplexing capability is particularly valuable when used in screening pooled CAR T libraries, as it enables a more comprehensive analysis of each CAR variant’s immune profile. In addition, evaluation of multiple cytokine markers simultaneously from a single cell through the nanovial platform allows the identification of individual CAR T cells with tailored cytokine signatures. These profiles can then be associated with improved persistence, anti-tumor activity, and potentially better safety outcomes in therapeutic settings in downstream characterization assays^35,48,49^. Future applications leveraging the polyfunctional screening offered by this platform could open up opportunities to refine CAR designs beyond basic activation markers and in addition to standard cytotoxicity cell killing assays, enabling the identification of constructs with more nuanced phenotypic profiles. This makes the nanovial platform not only useful for screening large libraries, but also for optimizing smaller, more rationally designed CAR T libraries.

Beyond using a pooled library of CAR constructs, the nanovial platform could unlock new functional insights from screening pooled libraries of genetic modulators, e.g., using CRISPR/Cas9 PERTURB-seq type approaches^50^. PERTURB-seq combines CRISPR-based screens with single-cell RNA-sequencing, enabling the systematic evaluation of gene functions. This would allow for a deeper understanding of the molecular signatures underlying the functional cytokine responses of CAR T cells, linking gene targets to specific functional phenotypes. By connecting genetic and functional data, we could identify key pathways or genetic drivers that contribute to optimal CAR T cell function, advancing the design of next-generation CAR constructs. Complementary linkages between genes and function may be obtainable by using SEC-seq approaches that link the amount of secreted cytokines, labeled with oligo-barcoded antibodies, with gene expression profiles at the single-cell level^25^.

The nanovial platform offers a highly versatile, scalable, and function-first approach for the analysis of CAR T cells and screening of pooled CAR T cell libraries. Unique features of the platform include the ability to quantitatively tune the level of target antigen and other features of the artificial immune synapse, such as inclusion of stimulatory or inhibitory co-receptors (e.g., 4-1BBL or PD-L1, respectively), enabling more mechanistic understanding through demultiplexing of the system. Compatibility with both FACS and single-cell sequencing allows enhanced scale of experiments - millions of cells screened - and molecular insights associated with functional responses. The scale of experiments and depth of data per individual cell could be used for generating perturbed data sets for training of AI models of cell function and cell-cell interactions for CAR T cells and other immune cell-tumor cell interactions, ultimately advancing the understanding and development of novel cancer therapies.

## Methods

### Nanovial Fabrication

Nanovials were fabricated using a flow-focusing microfluidic droplet generator as previously described^22^. Briefly, a PDMS-based flow-focusing droplet generator was used to co-flow the nanovial precursors, PEG solution and gelatin solution, with oil solution. PEG solution was made by dissolving 27.5% (w/v) 4-arm, 5kDa PEG-acrylate (Advanced BioChemicals) and 4% (w/v) lithium phenyl-2,4,6-trimethylbenzophosphinate (Sigma-Aldrich) in 1X phosphate buffered saline (PBS) (ThermoFisher Scientific). Gelatin solution was made by dissolving 20% (w/v) cold water fish skin gelatin (Sigma-Aldrich) in sterile MilliQ water. Oil solution was comprised of 0.5% (v/v) Pico-Surf (Sphere Fluidics) in Novec^TM^ 7500 Engineered Fluid (3M). The PEG, gelatin, and oil solutions were co-flowed into the PDMS device at 1, 1, and 10 μL/min respectively. Once phase separation had occurred, the PEG phase was crosslinked through UV exposure before particle collection. Upon collection, cured droplet emulsions containing nanovials were broken using a solution of 20% (v/v) perfluoro-1-octanol (PFO) (Sigma-Aldrich) in Novec^TM^ 7500. Excess oil was then removed by mixing hexane into the oil phase before removing both hexane and oil phase after centrifugation, this step was repeated twice. To remove remaining hexane, nanovials were washed by adding sterile 70% (v/v) ethanol before centrifugation and aspiration of supernatant. After, nanovials were incubated with 10mM Sulfo-NHS-Biotin (APExBIO) overnight at 4°C. Lastly, nanovials were sterilized using 70% ethanol, and stored at 4°C in Particle Wash Buffer (1X Dulbecco’s Phosphate Buffered Saline (Gibco^TM^), 0.05% Pluronic F-127 (Sigma), 0.5% Bovine Serum Albumin (Sigma-Aldrich), and 1X Antibiotic-Antimycotic (Gibco^TM^)) for long-term storage.

### Nanovial Functionalization

- *Streptavidin Conjugation* Nanovials, diluted to around 37,000 particles/mL in Particle Wash Buffer, were incubated with an equal volume of 200μg/mL streptavidin (ThermoFisher Scientific) using a tube rotator at room temperature for 30 minutes. Excess reagent was removed by spinning down the solution using a benchtop minifuge, followed by supernatant removal. To wash the nanovials, fresh Particle Wash Buffer was added to resuspend the nanovials, followed by centrifugation to remove the Particle Wash Buffer. This wash step was repeated twice.
- *Antibodies and Antigen Conjugation* Control aCD45-functionalized nanovials were formed by incubating nanovials at 37,000 particles/mL with 20μg/mL (130nM) of each biotinylated aCD45 (BioLegend) and biotinylated aIFNγ (R&D Systems). rHER2 nanovials, also at 37,000 particles/mL, were incubated with 20μg/mL (270nM) of biotinylated rHER2 protein (Sino Biological) and 20μg/mL (130nM) of biotinylated aIFNγ (R&D Systems). For concentration sweep experiments, the concentration of biotinylated aIFNγ was kept consistent while rHER2 was adjusted. Nanovials were incubated with antibodies overnight at 4°C. Excess reagent was removed and conjugated nanovials were washed as described in the previous section.

### Primary human T cell culture

Isolated peripheral blood mononuclear cells (PBMCs) from anonymous de-identified healthy donors were acquired through the UCLA Virology Core. T cells were immediately isolated using Dynabeads^TM^ Untouched^TM^ Human T Cells Kit (Invitrogen) and frozen in liquid nitrogen until use. Upon thawing, both T cells and CAR T cells were cultured in X-VIVO15 medium (Lonza) supplemented with 10% fetal bovine serum (FBS), 1X Antibiotic-Antimycotic (Gibco^TM^), and 100U/mL recombinant IL-2 (STEMCELL Technologies) for at least 24 hours at 37°C.

### Generation of CAR T cell library

The CAR library as well as the individual aHER2 CAR T cells were cloned as previously described^9^. Buffy Coats were obtained from healthy donors through the Blutspendezentrum Basel (University of Basel) from which T cells were isolated (STEMCELL Technologies) and stored in liquid nitrogen. Upon thawing, T cells were activated with Human T-Activator CD3/CD28 Dynabeads (Thermo Fisher Scientific) at a 1:0.75 cell/bead ratio and cultured in X-VIVO15 medium (Lonza) supplemented with 5% FBS, 50µM β-mercaptoethanol, 100µg/mL Primocin (Invivogen) and 100 IU/mL IL-2 (PeproTech).

Engineered CAR T cells were generated with CRISPR/Cas9 by integrating the respective CAR gene into the *TRAC* locus. For this, HDR templates of the CAR constructs were amplified as previously described^8^. Two days after T cell activation, Dynabeads were removed and used for subsequent electroporation using the 4D-Nucleofector^TM^ System (Lonza). One million cells were washed with FACS buffer (1x PBS, 2% FBS and 1mM EDTA) and resuspended in 20µL P3 nucleofection buffer (Lonza) including 1µL of Cas9 RNP transfection mix (100µM gRNA, IDT) and 500ng DNA HDR template. Next, T cells were shocked with the EH-115 program. Immediately after the shock, 80µL of T cell culture medium were added to the Lonza cuvettes and incubated for 20min at 37°C for recovery. Then, 1µL/mL of Alt-R HDR Enhancer V2 (IDT) and 100 IU/mL IL2 in T cell media were added to the rested cells to a final volume of 200µL and incubated for another 5h. After the 5h, media was exchanged to remove the Alt-R HDR Enhancer V2. In total eight to ten million cells were shocked for the CAR library, and four to six million cells for the individual aHER2 CAR T cells. Cells were expanded for seven to nine days in T cell medium before CAR T cell enrichment using FACS sorting. Transfected cells were analyzed and sorted as previously described^9^.

### Isolation of IFNγ-secreting CAR T cells in nanovials

Nanovials were functionalized with rHER2 antigens and aIFNγ antibody. Both control T cells and cells from the aHER2 CAR T cell library were thawed and recovered for 24 hours before being loaded into functionalized nanovials. After stimulation on nanovials, samples were stained with calcein AM (diluted 1:5000 in Particle Wash Buffer) and BrilliantViolet421-conjugated IFNγ detection antibody (5μL per 187,000 nanovials). Staining was done for 30 minutes at 37°C. Excess fluorescent dyes were removed and samples were washed with Particle Wash Buffer twice. Flow cytometry analysis and FACS-based cell sorting was done using the SONY SH800S Cell Sorter and FlowJo_v10.8.1.

### Sequencing and identification of CAR variants

Following sorting, genomic DNA (gDNA) from cells within nanovials and the starting cell library cells was extracted using QuickExtract DNA Extraction Solution (Lucigen) as per the manufacturer’s protocol. Extracted gDNA was used as the template for a 2-step nested PCR strategy. The first PCR reaction, using primers F1 and R1 (S. Table 2) amplified the CAR integration site within the TRAC locus to confirm CAR transgene insertion. PCR product was then purified using DNA Clean & Concentrator kit (Zymo Research). The purified PCR product was then used as template for the second PCR step using primer mixes F2 and R2 (S. Table 2). This was done to amplify the CAR 3’UTR region which contained the barcode sequence used to determine each library variant’s identity. Following a final DNA cleanup step, samples were sequenced using Amplicon Sequencing services from Genewiz. The obtained sequences were analyzed using the Biostrings and Tidyverse packages in R Studio. To identify IFNγ-secreting CAR T cell variants, around 400,000 reads from 300-500 sorted cells were assessed. Fold changes of the library after cytokine secretion were calculated for different incubation time points (3h, 12h). Additionally, CAR assignment rates, absolute counts and frequency were also determined.

## Acknowledgements

We would like to acknowledge the UCLA Jonsson Comprehensive Cancer Center for their flow cytometry services and UCLA/CFAR Virology Core Laboratory for providing primary human PBMCs. The work is partially supported by grants TLD1K132758: KUH-ART AND CA256084 from the National Institutes of Health and grant 2023-332384 from the Chan Zuckerberg Initiative Donor Advised Fund (CZI DAF), an advised fund of the Silicon Valley Community Foundation. Figures were created in https://BioRender.com.

