## Supplementary Information for "Screening of a pooled library of chimeric antigen receptor T cells based on secretory function"

Table 1 – Reagents

| Reagent Name | Company | Catalog # |
|---|---|---|
| *Nanovial Fabrication* | | |
| 4-arm Poly(ethylene glycol) acrylate (MW: 5kDa) | Advanced BioChemicals | 4AP0902 |
| Gelatin from cold water fish skin (solid) | Sigma-Aldrich | G7041 |
| Lithium phenyl-2,4,6-trimethylbenzoylphosphinate | Sigma-Aldrich | 900889 |
| Pico-Surf™ (5% (w/w) in Novec™ 7500) | Sphere Fluidics | C024 |
| Novec™ 7500 Engineered Fluid | 3M | 7100134816 |
| 1H,1H,2H,2H-Perfluoro-1-octanol | Sigma-Aldrich | 370533 |
| Sulfo-NHS-Biotin | ApexBio | A8001 |
| *Particle Wash Buffer* | | |
| Dulbecco's Phosphate Buffered Saline (1X) | Gibco™ | 14190144 |
| Pluronic® F-127 | Sigma-Aldrich | P2443 |
| Bovine Serum Albumin | Sigma-Aldrich | A7906 |
| Antibiotic-Antimycotic (100X) | Gibco™ | 15240062 |
| *Nanovial Functionalization* | | |
| Streptavidin | ThermoFisher Scientific | 434302 |
| Biotin anti-human CD45 Ab, 2D1 | BioLegend | 368534 |
| Human IFNg Biotinylated Antibody | Biotechne / R&D Systems | BAF285 |
| Human HER2/ErbB2 Protein (ECD), Biotinylated | Sino Biological | 10004-HCCH-B |
| aHER2/ErbB2 Antibody – biotin | Sino Biological | 10004-MM01-B |
| Recombinant Human IFNg Protein | R&D Systems | 285-IF-100 |
| *Cell Culture* | | |
| X-VIVO15 Serum-free Media | Lonza | 04-418Q |
| Human Recombinant IL-2, ACF | STEMCELL Technologies | 78145 |
| Dynabeads™ Untouched™ Human T Cells Kit | Invitrogen™ | 11344D |
| *Flow Cytometry Stain* | | |
| Invitrogen Molecular Probes Calcein, AM | Fisher Scientific | C3099 |
| aCD340 (erbB2/HER2) ms mAb - PE/Cy7 | BioLegend | 324413 |
| aIFNg ms mAb - BV421 | BioLegend | 502532 |
| *gDNA Isolation and Processing* | | |
| QuickExtract DNA Extraction Solution | LGC Biosearch Technologies | QE0905T |
| DNA Clean & Concentrator - 5 | Zymo Research | D4013 |
| Q5® Hot Start High-Fidelity 2X Master Mix | New England Biolabs | M0494 |

Table 2 – Primer Sequences

| ID | Sequence (5' → 3') |
|---|---|
| F1 | GGTCAGACAAGCTCCCGGAAAAGGA |
| R1 | AGGTGTCCCTTCCCTGCTT |
| F2 mix | GTCACCTAAATGCTAGAGCTCGC |
| | aGTCACCTAAATGCTAGAGCTCGC |
| | tcGTCACCTAAATGCTAGAGCTCGC |
| R2 mix | TACACGGCATGCCTGCTATTCT |
| | aTACACGGCATGCCTGCTATTCT |
| | tcTACACGGCATGCCTGCTATTCT |

Table 3 – Library Barcodes

'NA' indicates the lack of domain in the listed location.

| Variant | ICD 1 | ICD 2 | CD3Z | Barcode |
|---|---|---|---|---|
| 41BB-41BB-Z | 41BB | 41BB | Z | ACGGCGTTTCA |
| 41BB-CD28-Z | 41BB | CD28 | Z | GCCGACCTATA |
| 41BB-CD40-Z | 41BB | CD40 | Z | CCCGTCATCGG |
| 41BB-CTLA4-Z | 41BB | CTLA4 | Z | ATGGTCTTAAC |
| 41BB-IL15RA-Z | 41BB | IL15RA | Z | ATCGCACTGTA |
| 41BB-NA-Z | 41BB | NA | Z | GCCGGTCTAGT |
| CD28-41BB-Z | CD28 | 41BB | Z | CTCGGCCTAAC |
| CD28-CD28-Z | CD28 | CD28 | Z | AATGATATTAC |
| CD28-CD40-Z | CD28 | CD40 | Z | CCCGCAATCCG |
| CD28-CTLA4-Z | CD28 | CTLA4 | Z | AAGGAATTCTA |
| CD28-IL15RA-Z | CD28 | IL15RA | Z | GACGTTATAAA |
| CD28-NA-Z | CD28 | NA | Z | CCAGCTGTACT |
| CD40-41BB-Z | CD40 | 41BB | Z | CAAGTCCTGAG |
| CD40-CD28-Z | CD40 | CD28 | Z | GCAGGTCTGAC |
| CD40-CD40-Z | CD40 | CD40 | Z | GCCGCCATGCC |
| CD40-CTLA4-Z | CD40 | CTLA4 | Z | CCAGCCGTACA |
| CD40-IL15RA-Z | CD40 | IL15RA | Z | CGGGCAGTGCG |
| CD40-NA-Z | CD40 | NA | Z | TACGCTATTAA |
| CTLA4-41BB-Z | CTLA4 | 41BB | Z | AAGGATATTAG |
| CTLA4-CD28-Z | CTLA4 | CD28 | Z | GACGGTGTTAG |
| CTLA4-CD40-Z | CTLA4 | CD40 | Z | CCTGGCGTACG |
| CTLA4-CTLA4-Z | CTLA4 | CTLA4 | Z | AGTGGGGTTCA |
| CTLA4-IL15RA-Z | CTLA4 | IL15RA | Z | TCTGCGTTTCC |
| CTLA4-NA-Z | CTLA4 | NA | Z | GACGATATACG |
| IL15RA-41BB-Z | IL15RA | 41BB | Z | CTCGAAATGCA |
| IL15RA-CD28-Z | IL15RA | CD28 | Z | ATAGAAATCCC |
| IL15RA-CD40-Z | IL15RA | CD40 | Z | TAGGTAATGGA |
| IL15RA-CTLA4-Z | IL15RA | CTLA4 | Z | GCCGCGATCCA |
| IL15RA-IL15RA-Z | IL15RA | IL15RA | Z | ATCGAATTGTT |
| IL15RA-NA-Z | IL15RA | NA | Z | ACGGGTATACG |
| 1st Gen (NA-NA-Z) | NA | NA | Z | AAAGTGTTTAA |
| NS-CAR | NA | NA | NA | CCGGCACTATC |

Table 4 – CAR Assignment

| 3h - 1 | **Total Reads** | **CAR Reads** | **CAR Assignment (%)** |
|---|---|---|---|
| Initial Library | 421214 | 385781 | 98.30629295 |
| Non-Secretors | 399604 | 370900 | 97.83418711 |
| IFNγ-Secretors | 410656 | 376466 | 98.63865528 |

| 3h - 2 | **Total Reads** | **CAR Reads** | **CAR Assignment (%)** |
|---|---|---|---|
| Initial Library | 303102 | 278559 | 98.80815195 |
| Non-Secretors | 387811 | 360742 | 98.84820731 |
| IFNγ-Secretors | 407643 | 382439 | 99.20091832 |

Table 5 – Raw Sequencing Results (3h)

| Variant | Absolute Count | | | Percentage (%) | | |
|---|---|---|---|---|---|---|
| | Initial Library | Non-Secretors | IFNγ-Secretors | Initial Library | Non-Secretors | IFNγ-Secretors |
| 41BB-41BB-Z | 4700 | 704 | 11 | 1.24 | 0.19 | 0.00 |
| 41BB-CD28-Z | 5720 | 10332 | 2139 | 1.51 | 2.85 | 0.58 |
| 41BB-CD40-Z | 14511 | 2160 | 20 | 3.83 | 0.60 | 0.01 |
| 41BB-CTLA4-Z | 7035 | 12865 | 165 | 1.85 | 3.55 | 0.04 |
| 41BB-IL15RA-Z | 11727 | 6293 | 30 | 3.09 | 1.73 | 0.01 |
| 41BB-NA-Z | 5463 | 1433 | 92 | 1.44 | 0.39 | 0.02 |
| CD28-41BB-Z | 1275 | 3338 | 1426 | 0.34 | 0.92 | 0.38 |
| CD28-CD28-Z | 4419 | 3200 | 8784 | 1.17 | 0.88 | 2.37 |
| CD28-CD40-Z | 5371 | 11192 | 2705 | 1.42 | 3.08 | 0.73 |
| CD28-CTLA4-Z | 2079 | 1085 | 7 | 0.55 | 0.30 | 0.00 |
| CD28-IL15RA-Z | 4976 | 10050 | 10438 | 1.31 | 2.77 | 2.81 |
| CD28-NA-Z | 9982 | 8435 | 30631 | 2.63 | 2.32 | 8.25 |
| CD40-41BB-Z | 16971 | 21777 | 3928 | 4.47 | 6.00 | 1.06 |
| CD40-CD28-Z | 1812 | 4663 | 2585 | 0.48 | 1.29 | 0.70 |
| CD40-CD40-Z | 4330 | 1742 | 1543 | 1.14 | 0.48 | 0.42 |
| CD40-CTLA4-Z | 3855 | 1831 | 1350 | 1.02 | 0.50 | 0.36 |
| CD40-IL15RA-Z | 6644 | 9758 | 1709 | 1.75 | 2.69 | 0.46 |
| CD40-NA-Z | 8539 | 18032 | 14100 | 2.25 | 4.97 | 3.80 |
| CTLA4-41BB-Z | 1085 | 1544 | 342 | 0.29 | 0.43 | 0.09 |
| CTLA4-CD28-Z | 1117 | 527 | 1 | 0.29 | 0.15 | 0.00 |
| CTLA4-CD40-Z | 961 | 94 | 3 | 0.25 | 0.03 | 0.00 |
| CTLA4-CTLA4-Z | 647 | 355 | 1 | 0.17 | 0.10 | 0.00 |
| CTLA4-IL15RA-Z | 7171 | 1114 | 72761 | 1.89 | 0.31 | 19.59 |
| CTLA4-NA-Z | 720 | 37 | 1 | 0.19 | 0.01 | 0.00 |
| IL15RA-41BB-Z | 25105 | 6224 | 14276 | 6.62 | 1.72 | 3.84 |
| IL15RA-CD28-Z | 5165 | 5859 | 12142 | 1.36 | 1.61 | 3.27 |
| IL15RA-CD40-Z | 8651 | 7607 | 7736 | 2.28 | 2.10 | 2.08 |
| IL15RA-CTLA4-Z | 3745 | 2309 | 8 | 0.99 | 0.64 | 0.00 |
| IL15RA-IL15RA-Z | 25471 | 25144 | 113460 | 6.72 | 6.93 | 30.55 |
| IL15RA-NA-Z | 6949 | 3144 | 5598 | 1.83 | 0.87 | 1.51 |
| NA-NA-Z | 102749 | 161091 | 63206 | 27.09 | 44.39 | 17.02 |
| NA-NA-NA | 70302 | 18928 | 143 | 18.54 | 5.22 | 0.04 |

Table 6 – Raw Sequencing Results (3h-2)

| Variant | Absolute Count | | | Percentage (%) | | |
|---|---|---|---|---|---|---|
| | Initial Library | Non-Secretors | IFNγ-Secretors | Initial Library | Non-Secretors | IFNγ-Secretors |
| 41BB-41BB-Z | 514 | 400 | 1 | 0.19 | 0.11 | 0.00 |
| 41BB-CD28-Z | 849 | 1986 | 969 | 0.31 | 0.56 | 0.26 |
| 41BB-CD40-Z | 144 | 337 | 0 | 0.05 | 0.09 | 0.00 |
| 41BB-CTLA4-Z | 123 | 0 | 0 | 0.04 | 0.00 | 0.00 |
| 41BB-IL15RA-Z | 205 | 134 | 0 | 0.07 | 0.04 | 0.00 |
| 41BB-NA-Z | 473 | 105 | 0 | 0.17 | 0.03 | 0.00 |
| CD28-41BB-Z | 97 | 35 | 0 | 0.04 | 0.01 | 0.00 |
| CD28-CD28-Z | 224 | 461 | 0 | 0.08 | 0.13 | 0.00 |
| CD28-CD40-Z | 309 | 54 | 0 | 0.11 | 0.02 | 0.00 |
| CD28-CTLA4-Z | 120 | 0 | 0 | 0.04 | 0.00 | 0.00 |
| CD28-IL15RA-Z | 139947 | 203706 | 44770 | 50.85 | 57.13 | 11.80 |
| CD28-NA-Z | 69416 | 88716 | 76320 | 25.22 | 24.88 | 20.12 |
| CD40-41BB-Z | 1040 | 1459 | 1 | 0.38 | 0.41 | 0.00 |
| CD40-CD28-Z | 483 | 591 | 0 | 0.18 | 0.17 | 0.00 |
| CD40-CD40-Z | 13786 | 13252 | 1569 | 5.01 | 3.72 | 0.41 |
| CD40-CTLA4-Z | 146 | 233 | 0 | 0.05 | 0.07 | 0.00 |
| CD40-IL15RA-Z | 1020 | 803 | 0 | 0.37 | 0.23 | 0.00 |
| CD40-NA-Z | 25898 | 30277 | 145531 | 9.41 | 8.49 | 38.36 |
| CTLA4-41BB-Z | 208 | 241 | 0 | 0.08 | 0.07 | 0.00 |
| CTLA4-CD28-Z | 1232 | 837 | 1575 | 0.45 | 0.23 | 0.42 |
| CTLA4-CD40-Z | 57 | 321 | 0 | 0.02 | 0.09 | 0.00 |
| CTLA4-CTLA4-Z | 110 | 219 | 0 | 0.04 | 0.06 | 0.00 |
| CTLA4-IL15RA-Z | 119 | 0 | 0 | 0.04 | 0.00 | 0.00 |
| CTLA4-NA-Z | 120 | 188 | 0 | 0.04 | 0.05 | 0.00 |
| IL15RA-41BB-Z | 1872 | 2332 | 1 | 0.68 | 0.65 | 0.00 |
| IL15RA-CD28-Z | 6522 | 4898 | 98381 | 2.37 | 1.37 | 25.93 |
| IL15RA-CD40-Z | 3139 | 32 | 0 | 1.14 | 0.01 | 0.00 |
| IL15RA-CTLA4-Z | 236 | 0 | 4278 | 0.09 | 0.00 | 1.13 |
| IL15RA-IL15RA-Z | 510 | 579 | 0 | 0.19 | 0.16 | 0.00 |
| IL15RA-NA-Z | 2687 | 1219 | 2018 | 0.98 | 0.34 | 0.53 |
| NA-NA-Z | 1831 | 2304 | 3969 | 0.67 | 0.65 | 1.05 |
| NA-NA-NA | 1802 | 868 | 0 | 0.65 | 0.24 | 0.00 |

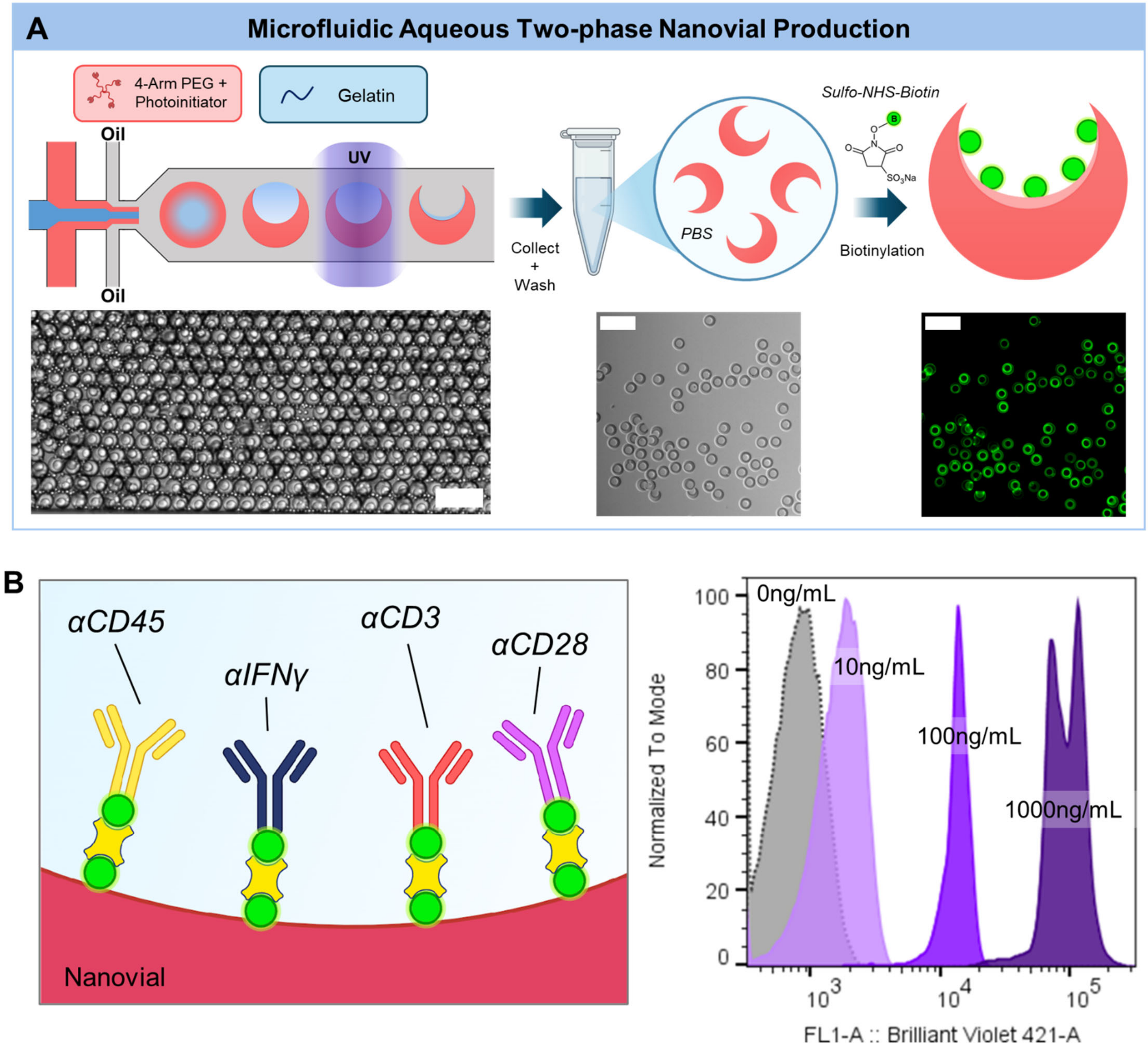


**Supplementary Figure 1**. Nanovial manufacture and cytokine detection dynamic range. (A) Schematic of nanovial fabrication process. Polyethylene glycol (PEG) and gelatin solution are co-flowed into a microfluidic device and pinched using oil to create droplets. PEG and gelatin solution phase-separate in parallel channels before UV curing of the PEG phase. Nanovials are then collected, washed, and stored in PBS-based solution (Particle Wash Buffer). The surface of the nanovial cavity is biotinylated using sulfo-NHS-biotin. Fluorescence image shows final nanovial product stained with Streptavidin-AlexaFluor488. Scale bar: 100µm. (B) Dynamic range of IFNγ capture signal on nanovials decorated with 4 different antibodies: anti-CD45, anti-CD3, anti-CD28, and anti-IFNγ.

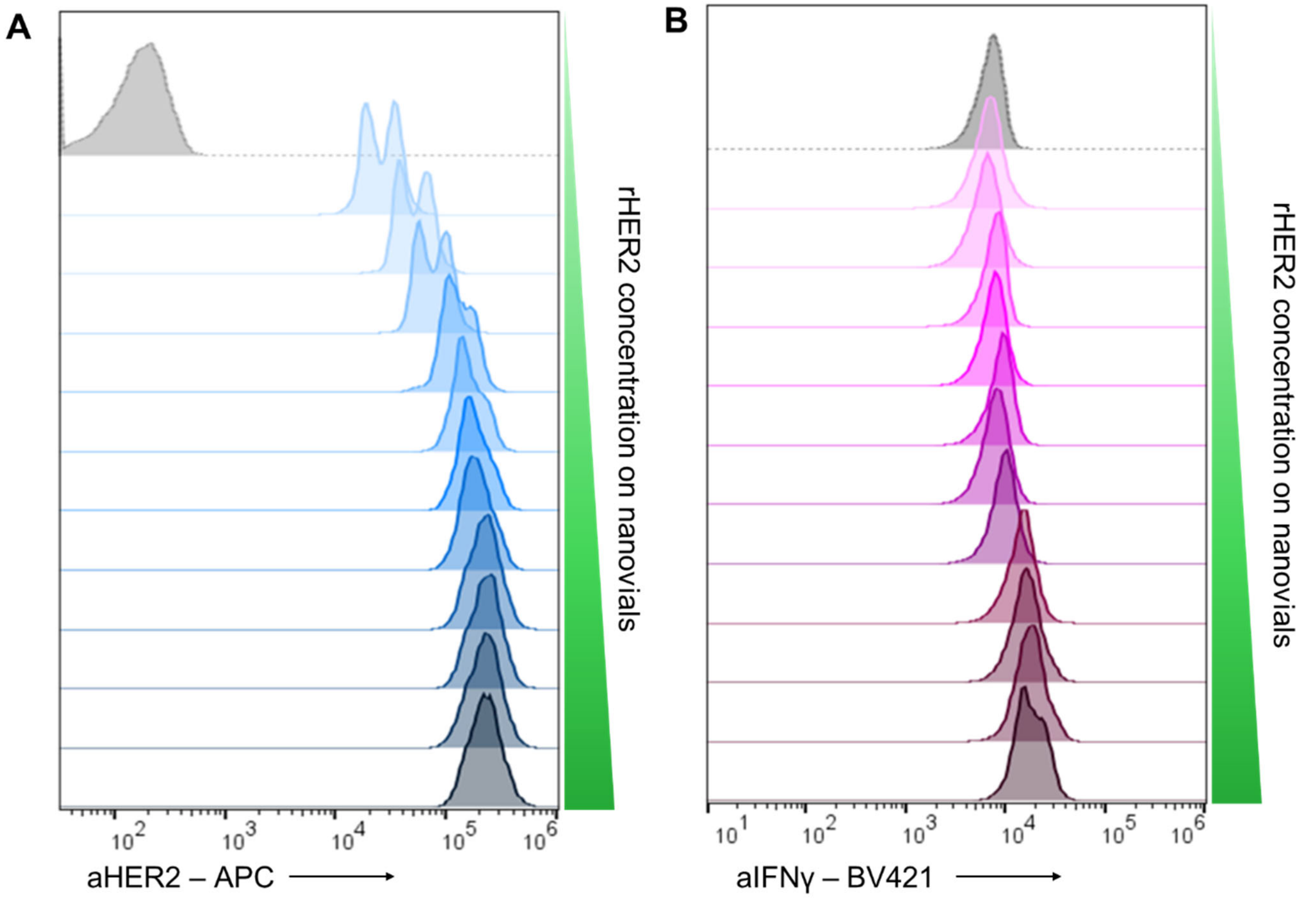


**Supplementary Figure 2.** Flow cytometry histograms of nanovials incubated with 50ng/mL recombinant IFNγ and increasing rHER2 concentrations. (A) Fluorescence stain of detected rHER2 immobilized on nanovials. (B) Fluorescence stain of detected recombinant IFNγ on nanovials.

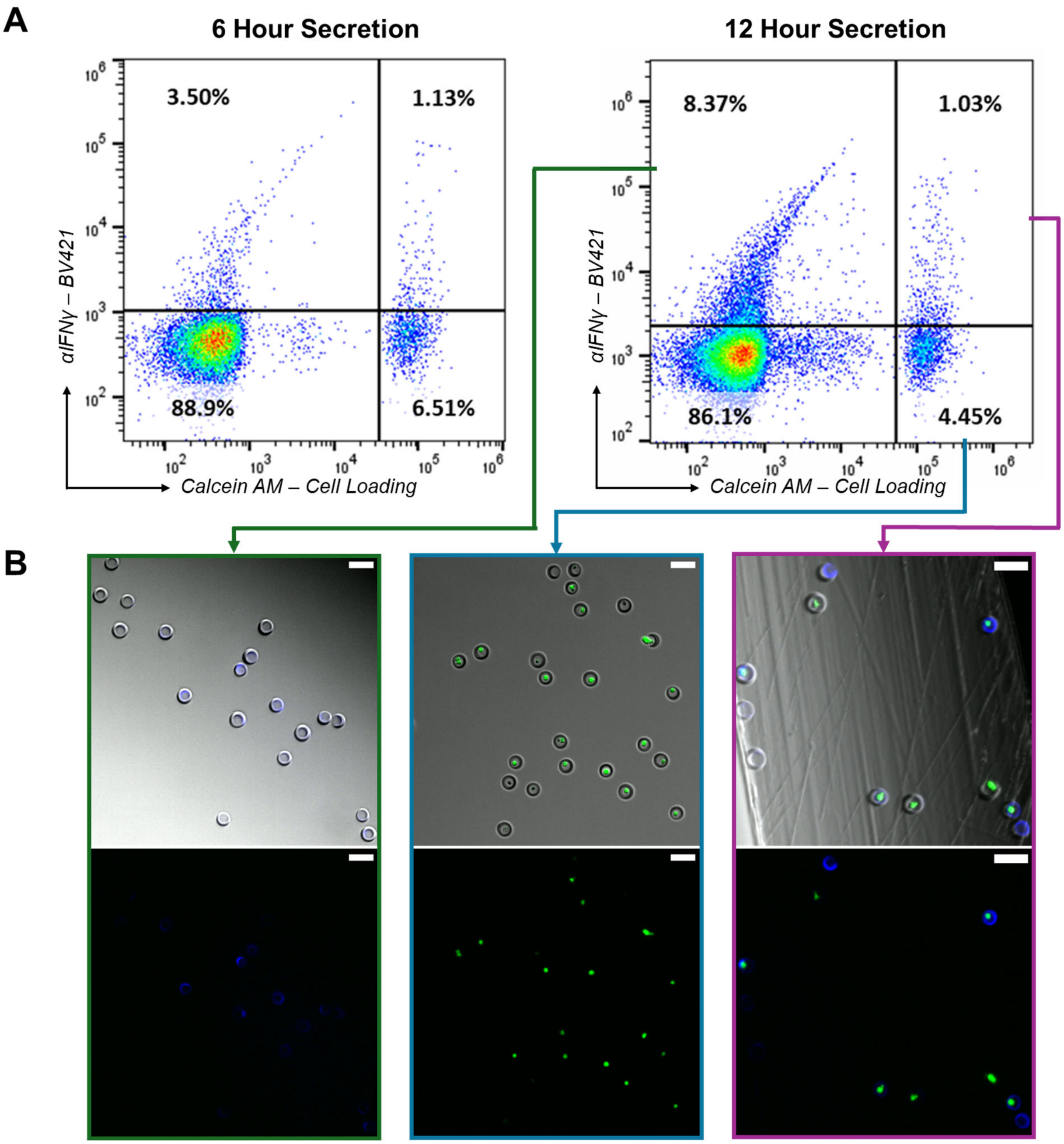


**Supplementary Figure 3.** Time-dependent secretion of IFNγ by aHER2 CAR T cells. (A) Flow scatter plots of calcein AM staining vs. IFNγ signal for aHER2 CAR T cells interacting with rHER2 functionalized nanovials for 6 and 12 hours. Gated quadrants define live cell-loaded versus unloaded nanovials and nanovials with secretion signal above threshold or no secretion signal. (B) Fluorescence microscopy images of gated populations. Green signal corresponds to calcein AM signal on cells and blue signal corresponds to IFNγ secretion on nanovials. Scale bar: 100μm

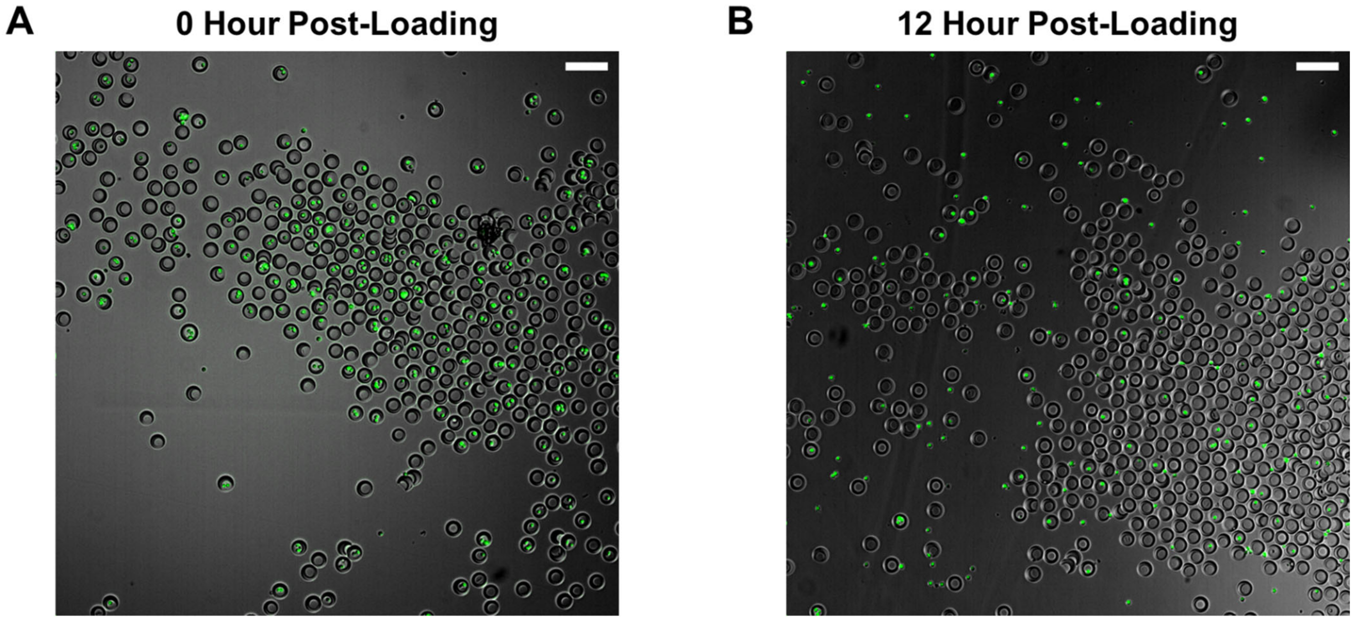


**Supplementary Figure 4.** Cell migration from nanovials over time. Fluorescence microscopy images (A) immediately after aHER2 CAR T cell loading into rHER2-coated nanovials and (B) after 12 hours of incubation. Green signal corresponds to calcein AM labeled CAR T cells. More cells are seen outside of nanovials on the surrounding well surface at 12 hours. Scale bar: 100µm

A

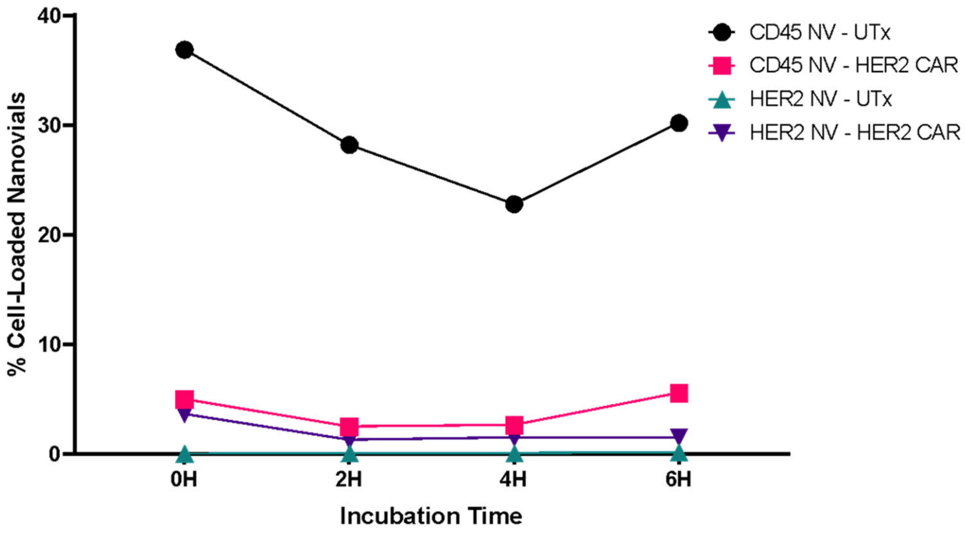


B

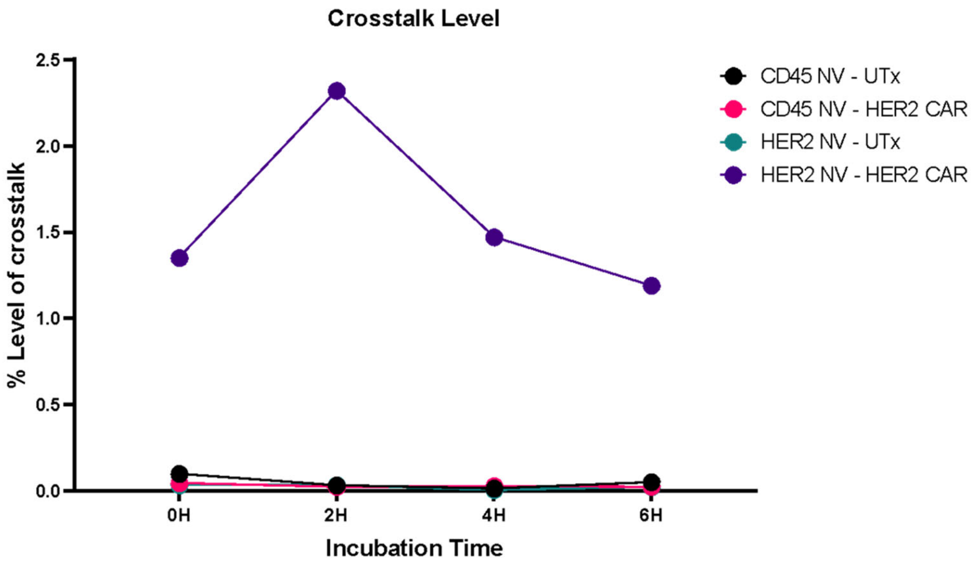


**Supplementary Figure 5.** Time dependence of CAR T cell loading and crosstalk on rHER2-functionalized nanovials. (A) Percentage of cell-loaded nanovials within 0-6 hours of incubation remain similar. (B) Level of crosstalk between nanovials also remains similar within 0-6 hours of incubation. Crosstalk is defined as nanovials without any cell loaded but show IFNγ capture signal, most likely due to spillover from neighboring nanovials.

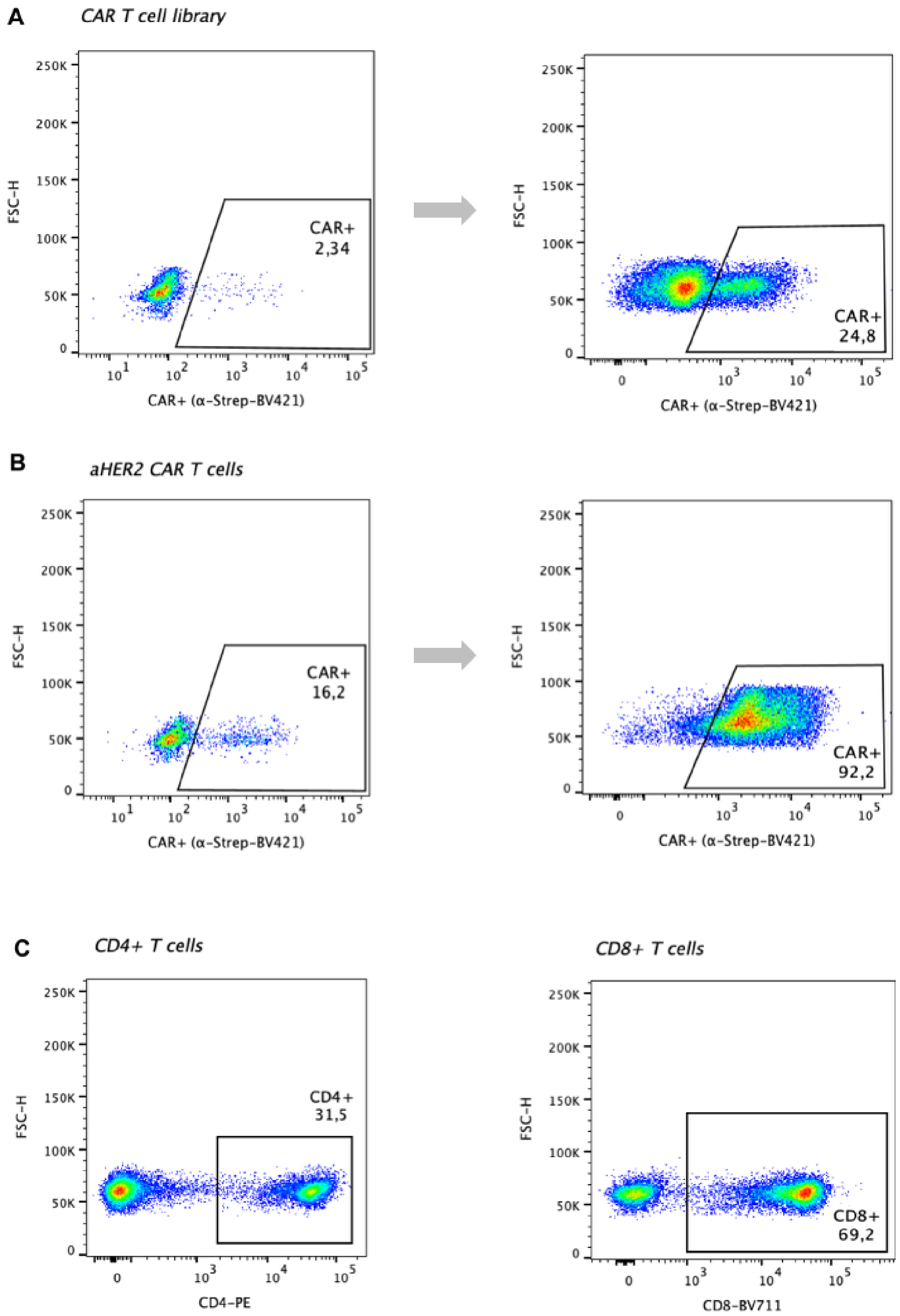


**Supplementary Figure 6.** FACS plots showing the enrichment and characterization of the CAR T cell library and the aHER2 CAR T cells. (A) Positive fraction of CARs in the library before and after a FACS sort. (B) Positive fraction of the aHER2 CARs before and after a FACS sort. (C) Distribution of CD4+ and CD8+ T cells in the CAR T cell library after enrichment.

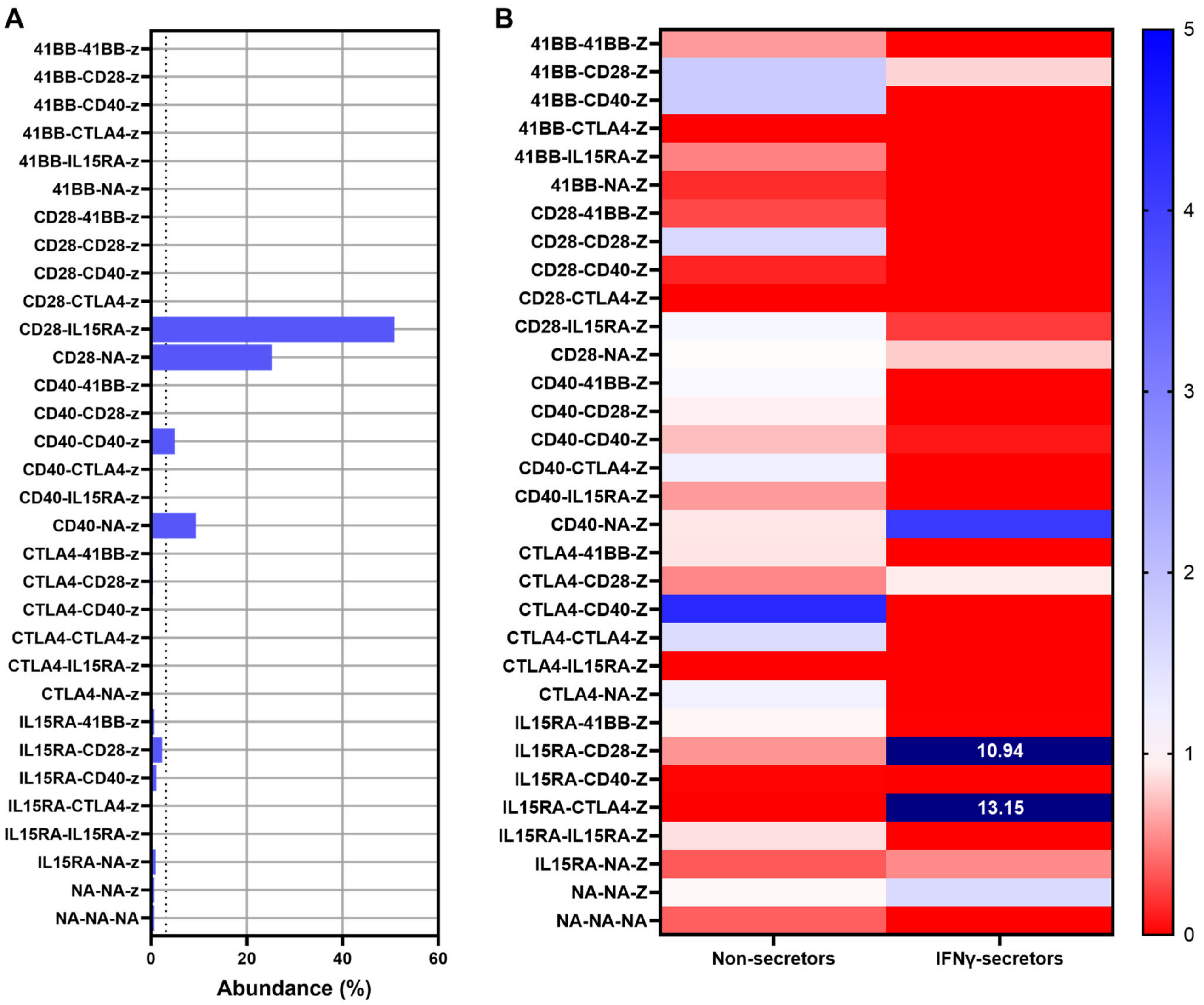


**Supplementary Figure 7.** (A) Initial distribution of the aHER2 CAR T cell library used in the repeat 3-hour incubation screen. Dashed line represents a theoretical equal distribution of constructs of ~3%. (B) Heatmap showing relative depletion (values from 0-1) or enrichment (values >1) of constructs normalized to their respective starting library frequency for the repeated 3-hour incubation screen.

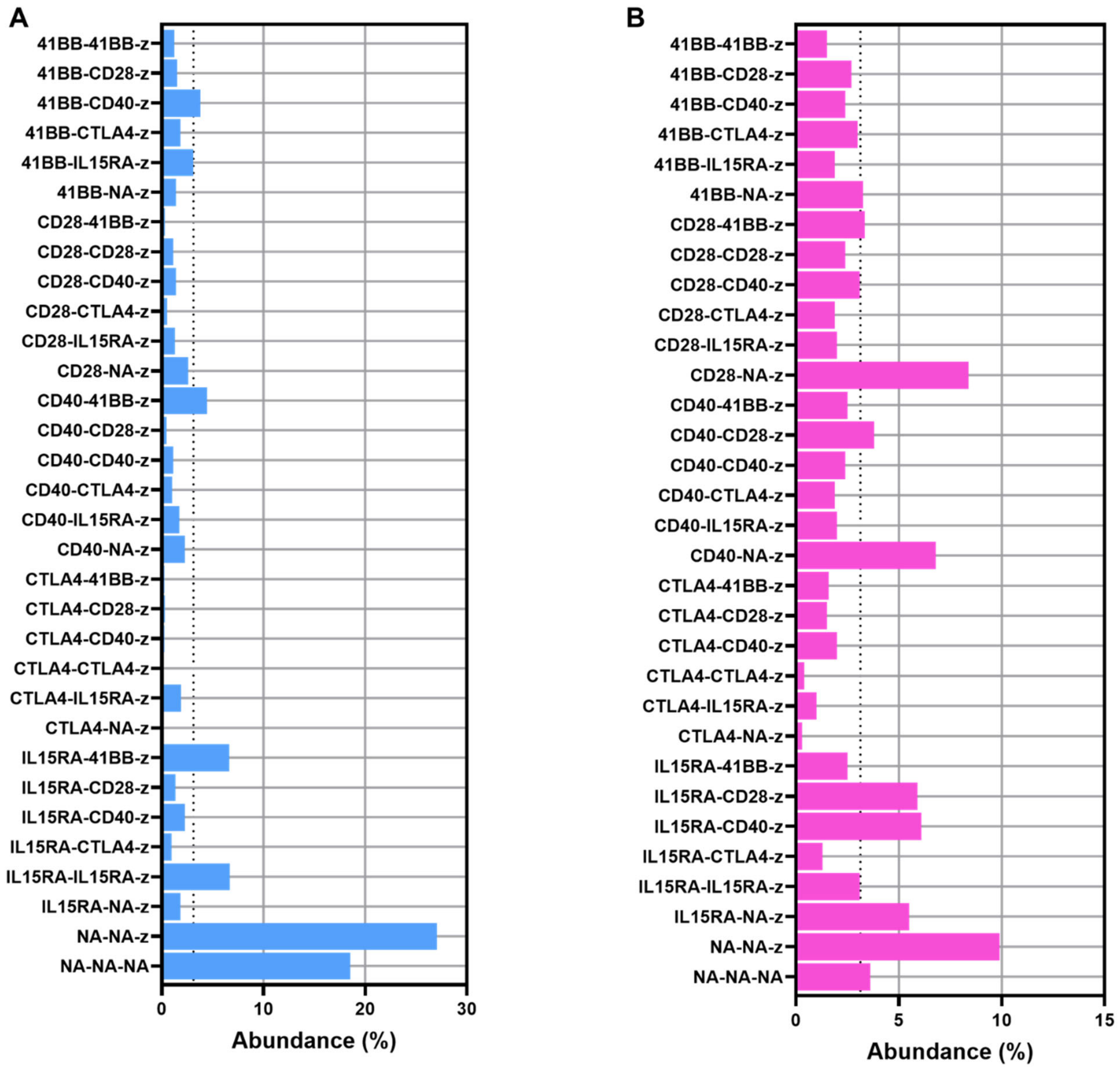


**Supplementary Figure 8.** (A) The CAR variant distribution of the starting aHER2 CAR T cell library for the first 3-hour incubation screen determined by DNA barcodes. Dashed line represents a theoretical equal distribution (~3%) of the CAR constructs. (B) As for **A** but for the aHER2 CAR T cell library used in the 12-hour incubation screen.
